# Leveraging high-throughput screening data and conditional generative adversarial networks to advance predictive toxicology

**DOI:** 10.1101/2020.10.02.322917

**Authors:** Adrian J. Green, Martin J. Mohlenkamp, Jhuma Das, Meenal Chaudhari, Lisa Truong, Robyn L. Tanguay, David M. Reif

## Abstract

There are currently 85,000 chemicals registered with the Environmental Protection Agency (EPA) under the Toxic Substances Control Act, but only a small fraction have measured toxicological data. To address this gap, high-throughput screening (HTS) methods are vital. As part of one such HTS effort, embryonic zebrafish were used to examine a suite of morphological and mortality endpoints at six concentrations from over 1,000 unique chemicals found in the ToxCast library (phase 1 and 2). We hypothesized that by using a conditional Generative Adversarial Network (cGAN) and leveraging this large set of toxicity data, plus chemical structure information, we could efficiently predict toxic outcomes of untested chemicals. CAS numbers for each chemical were used to generate textual files containing three-dimensional structural information for each chemical. Utilizing a novel method in this space, we converted the 3D structural information into a weighted set of points while retaining all information about the structure. *In vivo* toxicity and chemical data were used to train two neural network generators. The first used regression (Go-ZT) while the second utilized cGAN architecture (GAN-ZT) to train a generator to produce toxicity data. Our results showed that both Go-ZT and GAN-ZT models produce similar results, but the cGAN achieved a higher sensitivity (SE) value of 85.7% vs 71.4%. Conversely, Go-ZT attained higher specificity (SP), positive predictive value (PPV), and Kappa results of 67.3%, 23.4%, and 0.21 compared to 24.5%, 14.0%, and 0.03 for the cGAN, respectively. By combining both Go-ZT and GAN-ZT, our consensus model improved the SP, PPV, and Kappa, to 75.5%, 25.0%, and 0.211, respectively, resulting in an area under the receiver operating characteristic (AUROC) of 0.663. Considering their potential use as prescreening tools, these models could provide *in vivo* toxicity predictions and insight into untested areas of the chemical space to prioritize compounds for HT testing.

**Summary:** A conditional Generative Adversarial Network (cGAN) can leverage a large chemical set of experimental toxicity data plus chemical structure information to predict the toxicity of untested compounds.

## Introduction

Currently, there are 85,000 chemicals registered with the EPA, as part of the Toxic Substances Control Act[1], that are manufactured, processed, or imported into the United States; however, only 4,400 have rigorous toxicological data, leaving over 80,000 chemicals untested [2,3]. Due to the high cost, and ethical concerns over the use of low-throughput mammalian models associated with traditional *in vitro* and *in vivo* assays, there has been increasing demand to reduce the number of animals used in toxicity testing paradigms by switching to *in silico* methods [4]. To directly address this chemical data gap and help prioritize chemicals for testing, the ToxCast program was developed[5,6]. The original ToxCast program included approximately 700 biochemical and cell-based assays, which was efficient but lacking in systemic biological complexity[2]. Therefore, as part of an effort to expand the toxicology database, a multidimensional, high-throughput screening (HTS) assay was devised to examine all ToxCast phase 1 and 2 chemicals (over 1,000 unique chemicals) for developmental and neuro-toxicity in the embryonic zebrafish [7].

Additionally, computational approaches to bridge the data gap above have been employed, with Quantitative Structure-Activity Relationship (QSAR) and Read-Across being the most commonly used methodologies [8–13]. Both methods rely on the grouping of chemicals together using functional features, e.g. number of carbons, and have employed statistical or machine learning approaches. Although these methods have been useful in identifying priority compounds for further testing, how these chemicals are grouped together might add bias and recent machine learning advances have not been thoroughly explored [14]. Therefore, we propose using a supervised generative adversarial machine learning architecture that leverages the zebrafish HTS assay data along with chemical structure information to predict the toxic outcomes of untested chemicals. This approach will expand QSAR methodologies by providing a computational toxicology tool capable of prescreening the untested chemical space for active developmental toxicants.

## Materials and methods

In this section, we describe a new conditional generative adversarial network utilizing a novel weighted sum over views notation of compounds to predict active developmental toxicants. An overview of our approach is shown in Figs 1 and 2. First, we used experimental data collected on a large, diverse compound set to assess the toxic effects of these chemicals following developmental exposure (Fig 3a). Next, we recast the chemical data in a structural representation that maintained connectivity and positional information of each atom in the molecule but in a format easily read as input into a neural network. Next, we trained two types of generators to produce toxicity data using the recast chemical structural representation. The first used a cGAN architecture while the second utilized a deep neural network (DNN) with regression for training. This was done to produce generators capable of predicting developmental zebrafish toxicity data dependent on chemical structure alone. Regression training was leveraged to maximize model fit, minimize training time, and utilized a simpler toxicity data representation. cGAN training minimized the effects of outliers and increased network adaptability to chemical structure. These generators were trained using phase 1 and 2 ToxCast chemical data (n = 1003) split 80:20 into training and validation sets (Fig 3a). Finally, we evaluated the trained networks on an independent test set containing chemicals (n = 57) of greater diversity in terms of both size and atomic constituents (Fig 3b).

**Fig 1.**
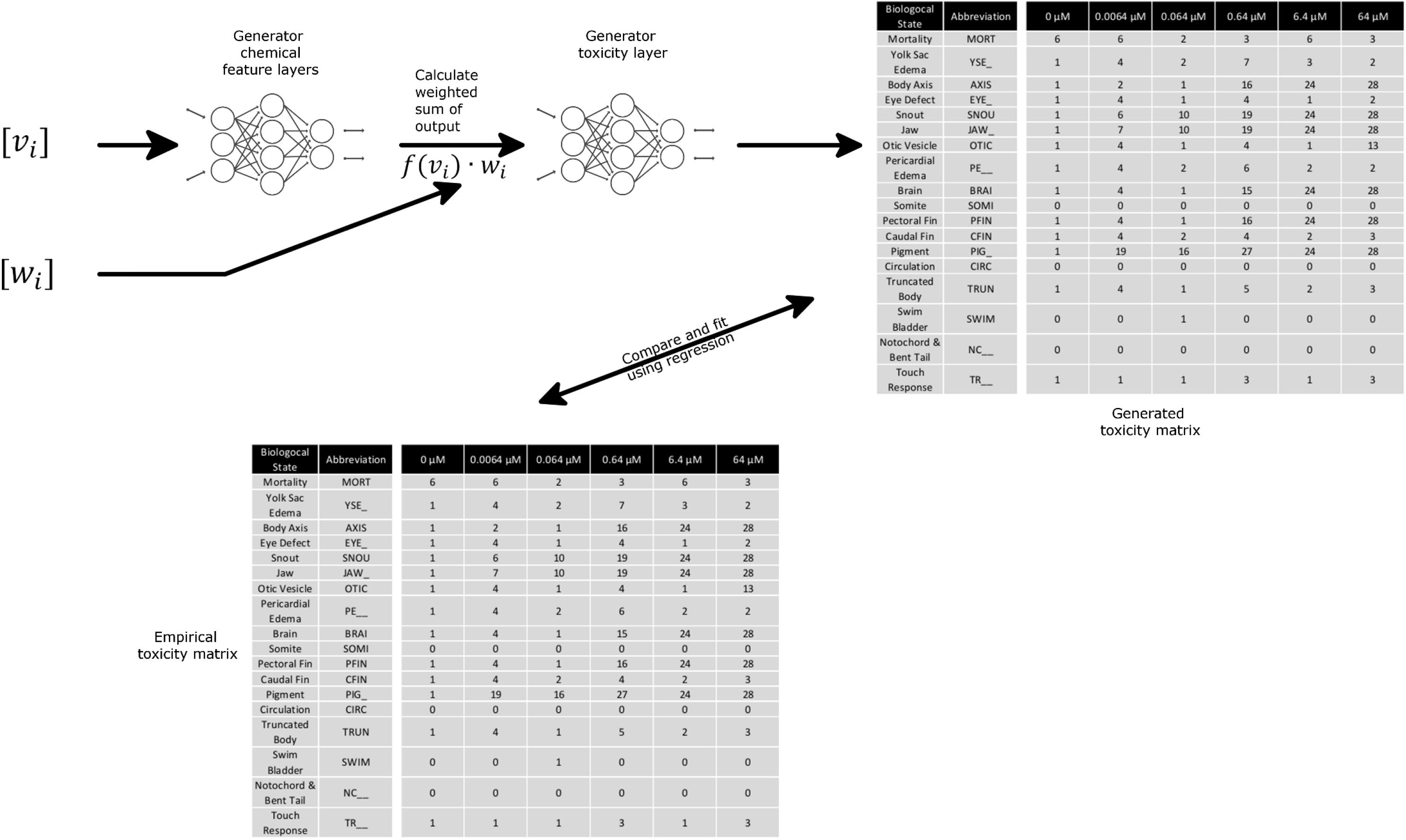
Regression generator diagram. Schematic representation of Go-ZT architecture showing chemical structural input represented as weights (w_*i*_) and views (v_*i*_) matrices passed through two fully connected neural networks to produce a predicted toxicity matrix.

**Fig 2.**
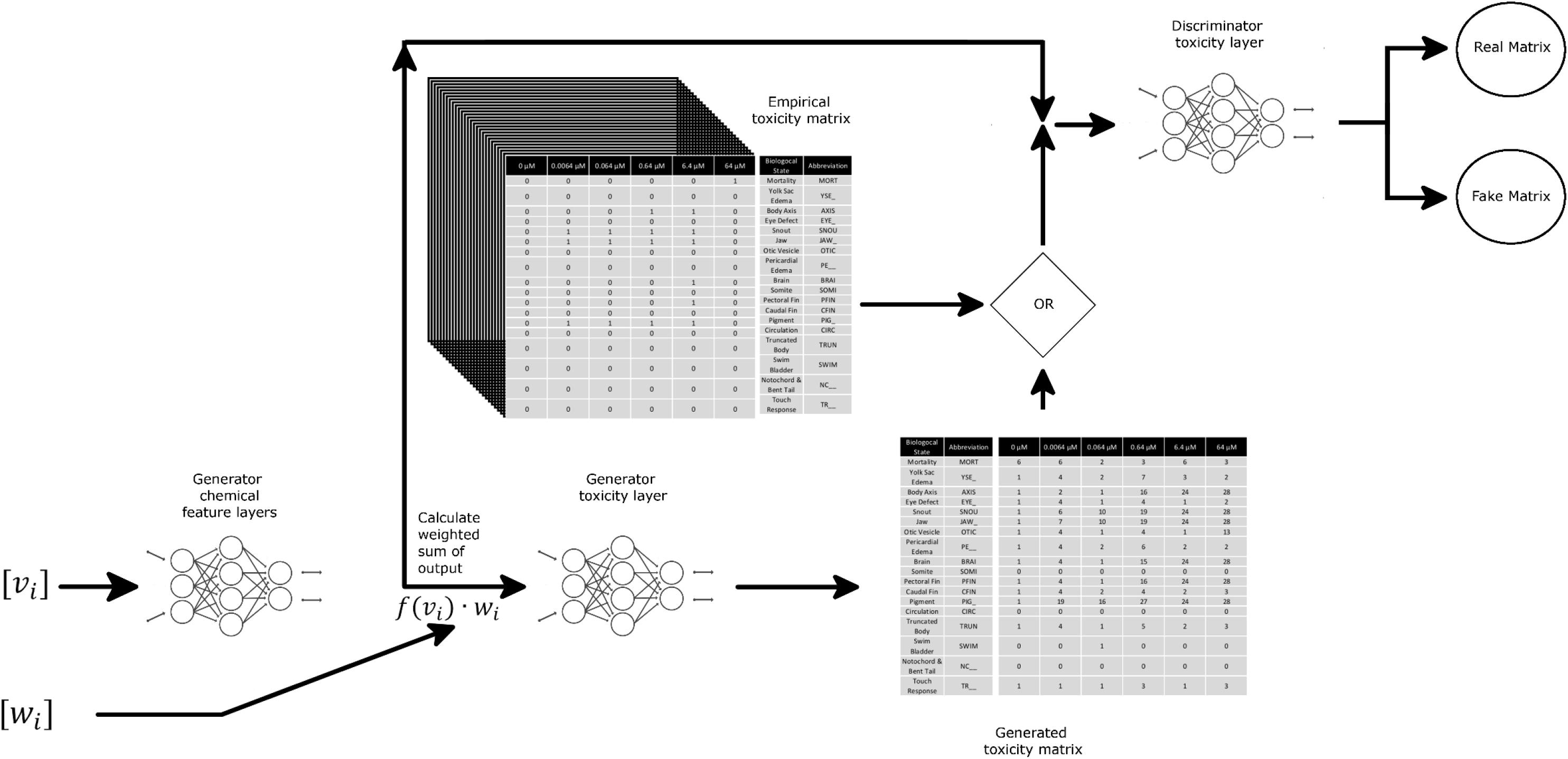
Conditional GAN diagram. Schematic representation of GAN-ZT architecture showing chemical structural input represented as weights (w_*i*_) and views (v_*i*_) matrices passed through two fully connected neural networks to produce a predicted toxicity matrix. Chemical features along with predicted or empirical toxicity matrices are then passed to a discriminator comprising a fully-connected neural network.

**Fig 3.**
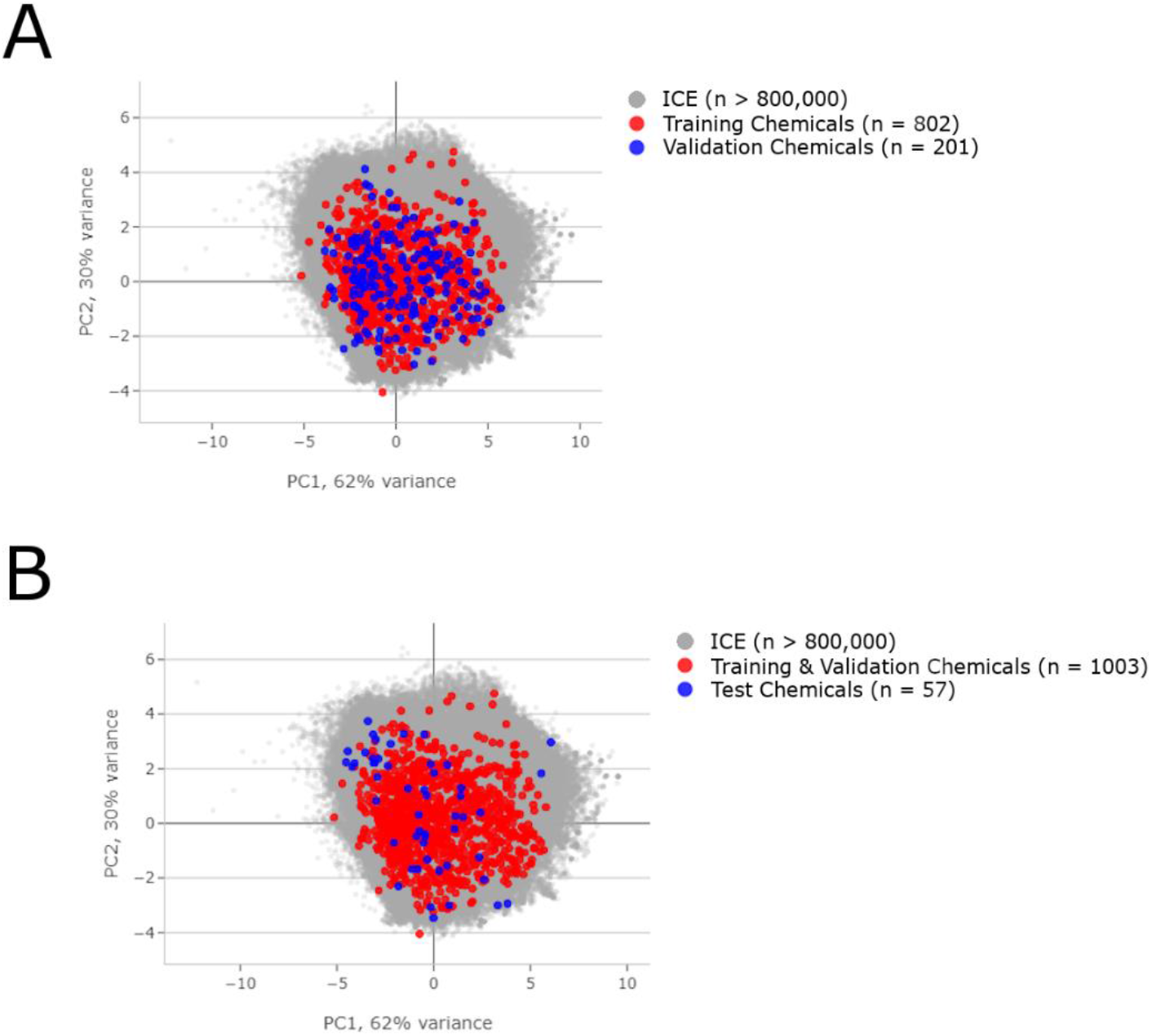
Data subdivision. Principal component analysis displayed against the background of over 800,000 chemicals in the EPA Integrated Chemical Environment database. (A) Compares physical chemical properties between the training and validation sets. (B) Compares physical chemical properties between the training and validation set, and the test set.

### Empirical (experimental) data

The empirical data used to develop a generator of zebrafish toxicity were gathered as described in Truong et al. and Noyes et al.[7,15]. Fig 4 shows the experimental design and input data structure. The data included 1003 unique ToxCast chemicals tested at six concentrations for each chemical (0 μM, 0.0064 μM,0.064 μM, 0.64 μM, 6.4 μM and 64 μM). To minimize effects of response variability at lower concentrations, only the highest concentration was chosen for network training. There were 32 replicates (an individual embryo in singular wells of a 96-well plate) at each concentration for each chemical. At 120 hours post-fertilization (hpf), 18 distinct developmental endpoints were evaluated. The data were recorded as binary incidences and split into training (n = 802) and validation (n = 201) chemical sets (Fig 3a). The data were either summed for each individual and normalized by the number of larvae per concentration or kept as individual 1 by 18 matrices and used to train, and validate our regression and GAN networks, respectively. In a similar manner, toxicity matrices were created for an independent test set of 57 chemicals that were collected in new experiments after collection of the original ToxCast data. This new test chemical set was more diverse in terms of atomic species and physical chemical properties (Fig 3b)[16]. Due to chemical vectorization constraints we defined a reasonable domain of applicability to exclude, Perfluorinated chemicals with carbon chains longer than nine, Chloroperfluoro chemicals, and chemicals with a betaine functional group. Fig 3 shows the division of these data into train/validate/test subsets and Principal Component Analysis (PCA) comparisons of the physical chemical properties using the EPA’s Integrated Chemical Environment chemical characterization tool[16]. The PCA analysis includes the following physical chemical properties: Molecular Weight, Boiling Point, Henry’s Law, Constant Melting Point, Negative Log of Acid Dissociation Constant, Octanol-Air Partition Coefficient, Octanol-Water Distribution Coefficient, Octanol-Water Partition Coefficient, Vapor Pressure, and Water Solubility[16].

**Fig 4.**
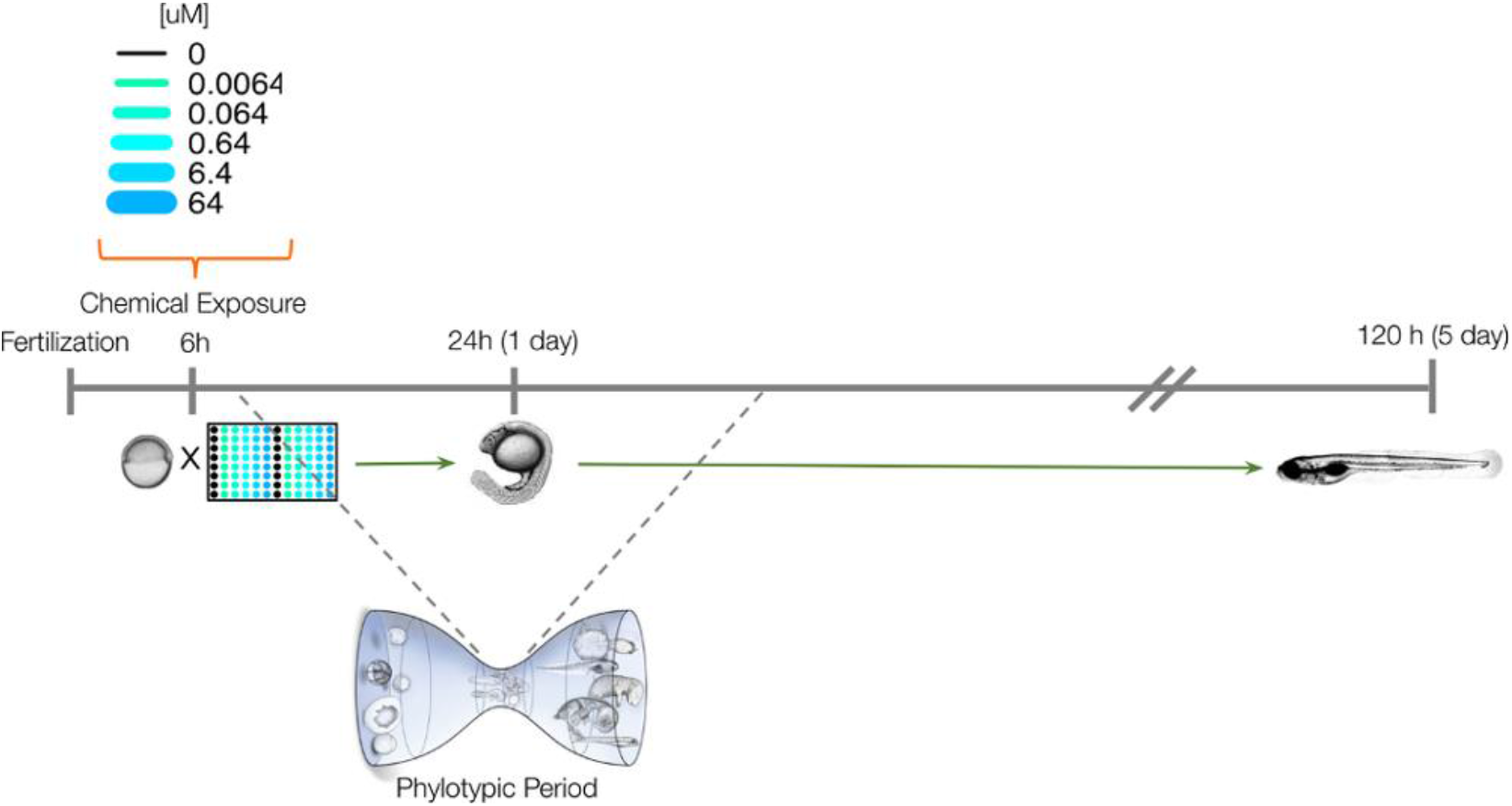
Experimental design. Schematic representation of the experimental approach for screening developmental and neurotoxicity of chemicals in larval zebrafish.

### Representing chemical compounds

Utilizing their CAS numbers chemical structural information was retrieved from the EPA’s Chemistry Dashboard and converted from SDF to PDB format using Open Babel[17,18]. The PDB format was chosen as it is easily accessible and contains 3D structural information for all atoms in a molecule. Multiple methods have been used to encode this structural information for utilization in deep learning, including, chemical properties, molecular fingerprints, SMILES, and graph vectorization, as well as 2D images of a chemical [19–21]. In this analysis, we utilized an algorithm developed to map and vectorize structure that was originally created for use in material sciences [22]. The PDB file for each chemical was vectorized as described by d’Avazac et al. [22] and illustrated in S1 Fig. This method is simple and universal with few parameters and was adapted as follows: First, each chemical was vectorized (x, y, and z coordinates) using either all atoms or only carbon atoms, when available, in the PDB list. The relative position of each additional atom in the PDB list was added to the view until all atoms were added or a user defined limit was reached. Lastly, an atom’s position on the periodic table was used as a unique identifier (group and period) producing five chemical features per atom.

### cGAN and regression generator

Two network architectures were developed and tested to train a generator that was capable of using 3D chemical information, in the form of vectorize views (S1 Fig), and generate a toxicity matrix (Figs 1 and 2). The following two different models were trained on a Dell R740 containing two Intel Xeon processors with 18 cores per processor, 512 GB RAM, and a Tesla-V100-PCIE (31.7 GB). The first was a simpler deep neural network trained to produce a toxicity matrix using regression (Fig 1). We used multiple layers consisting of a deep neural network base layer to extract salient features from the views matrix for each chemical structure (generator chemical feature layer). A second layer calculated the weighted sum over features (*f*(*v*_*i*_·*W*_*i*_) and a final deep neural network (generator toxicity layer) generates toxicity values. The regression generator (Go-ZT) was trained over the course of 200 epochs (35 seconds/run) and the generated toxicity matrices were compared to empirical toxicity matrices and the network that generated toxicity matrices with the highest Cohen’s Kappa statistic was used for further evaluation.

The second network developed (GAN-ZT) used a much more complex cGAN architecture to generate a toxicity matrix (Fig 2). Similar to the Go-ZT above, we used multiple layers consisting of a deep neural network base layer to extract salient features from the views matrix for each chemical structure (generator chemical feature layer) and a second layer to calculate the weighted sum over these features (*f*(*v*_*i*_)·*W*_*i*_). The resulting weighted sum of views (*f*(*v*_*i*_)·*W*_*i*_) for each chemical was used as input to a final deep neural network (generator toxicity layer) to produce a generated toxicity matrix. The discriminator took the generators resulting weighted sum of views (*f*(*v*_*i*_)·*W*_*i*_) for each chemical along with its corresponding empirical or generated toxicity matrices to determine whether the toxicity matrix was real or fake. This information was then backpropagated to train the generator. GAN-ZT was trained over the course of 500 epochs (1.5 hours/run) and the network that generated toxicity matrices with the highest Cohens Kappa statistic was used for further evaluation.

### Network performance and evaluation

Previous work analyzing this data has shown that a summary value, aggregate entropy (AggE) can be calculated from the 18 morphological endpoints[23]. The intent of this summary value is to capture a meaningful measure of toxicity, while avoiding overinflation by summing highly correlated endpoints. Using the threshold value (9.35) identified by Zhang et al. for AggE in these data, compounds may be classified as active or inactive[23]. It should be noted that this threshold value influences the toxicity hit rate of a chemical and is concentration dependent. Therefore, it would need to be changed when investigating other nominal concentrations. Following training, the resulting generators were used to output toxicity matrices for the 201 chemicals in the validation dataset. AggE values were calculated using both the empirical and generated toxicity matrices and compounds were classified as active or inactive using the threshold value identified above. Active vs inactive classification accuracy was evaluated using a confusion matrix, Cohen’s Kappa statistic, and area under the receiver operating characteristic (AUROC) as Kappa and AUROC measure model accuracy, while compensating for simple chance[24]. The primary metrics we used from the confusion matrix included sensitivity (SE), specificity (SP), and positive predictive value (PPV) as these parameters give us the true positive rate, true negative rate, and the proportion of true positives amongst all positive calls[25–27]. The network with the highest Kappa statistic and positive predictive value (PPV) was used for evaluation of the test dataset.

## Results

### Capacity ceiling and network training

The first step in training both Go-ZT and GAN-ZT included recasting the chemical structures as views and extracting relevant features from this chemical data. Therefore, at the very least our networks should be capable of extracting physical properties from the chemical data provided. As shown in S2 Fig, our GAN-ZT architecture generates chemical diameters with a Mean Absolute Percent Error (MAPE) of 12.4% and 13.0% for the training and validation datasets, respectively. While Go-ZT architecture generates chemical diameters with a MAPE of 10.3% and 13.8% for the training and validation datasets, respectively. Therefore, the best performance we could expect from either GAN-ZT or Go-ZT is approximately 13.0%.

Following training using chemical and toxicity data the best networks utilized Swish activation, batch normalization between layers, mean squared error for the loss function, and Adam as the optimizer[28–30]. GAN-ZT and Go-ZT generated toxicity values, using the chemical structures found in our training and validation datasets, with a validation MAPE of 49.1% and 51.6%, respectively (S3 Fig).

### Predicting toxicity

Empirical and generated toxicity matrices for each chemical from the validation or test datasets (Fig 3b) were used to calculate AggE values and determine activity classification. The empirical validation dataset contains 22 chemicals that meet the AggE threshold to be classified as active compounds (10.9%). As shown in Table 1, changing the number of chemical views from 43 (preferring views starting with Carbon) to 126 (maximum), decreasing the maximum number of atoms per view from 118 (maximum number of atoms in the training set) to 20 by truncation, and decreasing the number of chemical features from 590 (maximum) to 120 in training altered the SE, Kappa, and AUROC results of both the Go-ZT and GAN-ZT network architectures. GAN-ZT_v2 and Go-ZT_v3 were chosen for final evaluation using the test dataset as they had the best Kappa and AUROC values.

**Table 1.**
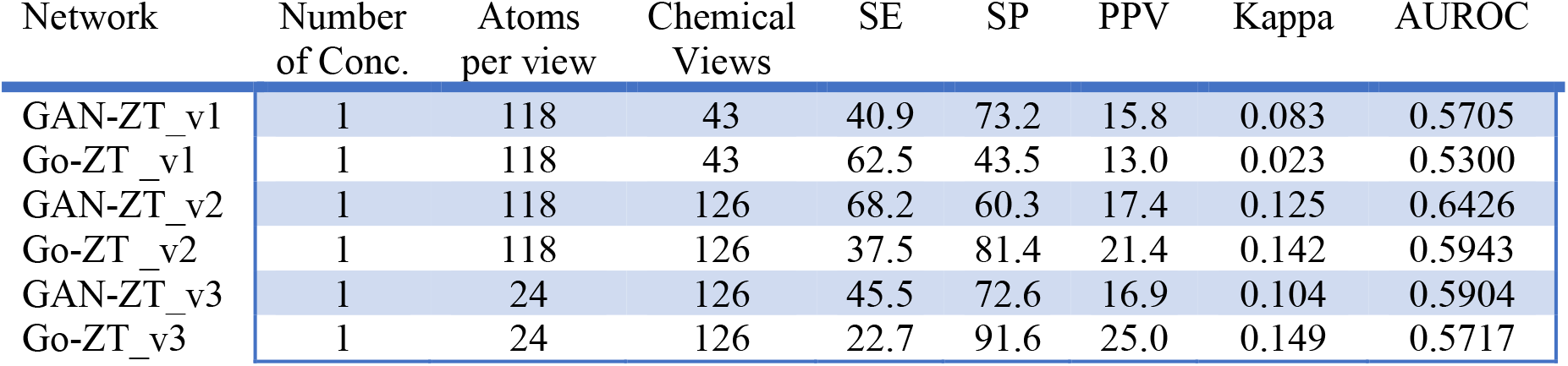
Prediction performances: validation chemicals.

The Go-ZT_v3 and GAN-ZT_v2 networks predicted 5 and 15 of the 22 active chemicals in the validation dataset, respectively. As shown in Table 1, this resulted in a SE, SP, and PPV for Go-ZT of 22.7%, 91.6%, and 25.0% respectively. GAN-ZT on the other hand produced SE, SP, and PPV values of 68.2%, 60.3%, and 17.4%, respectively. Evaluation of the chemical domain space using Go-ZT and GAN-ZT showed that the chemicals excluded due to long chain lengths, or betaine or Chloroperfluoro functional groups showed that these chemical properties fall outside of the domain space of our models.

Go-ZT and GAN-ZT were then used to predict the toxicity of an independent and more diverse chemical set containing seven active compounds (12.5%). The results (Table 2) show that Go-ZT performed best with increases in both Kappa and AUROC values while GAN-ZT saw declines in their Kappa and AUROC values. Overall Go-ZT had higher ACC, SP, and PPV while GAN-ZT had a better SE (Fig 5). By combining the predictive results of these two machine learning architectures we were able to leverage the sensitivity of the cGAN and the specificity of the regression model (Fig 6). As a result of the consensus between the models we were able to capture four of the seven active chemicals in the test set while eliminating 29 false positives which translates to a PPV of 25.0%.

**Table 2.**
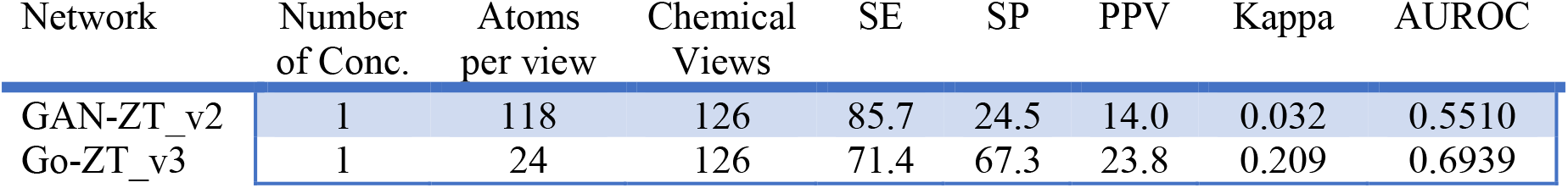
Prediction performances: test chemicals.

**Fig 5.**
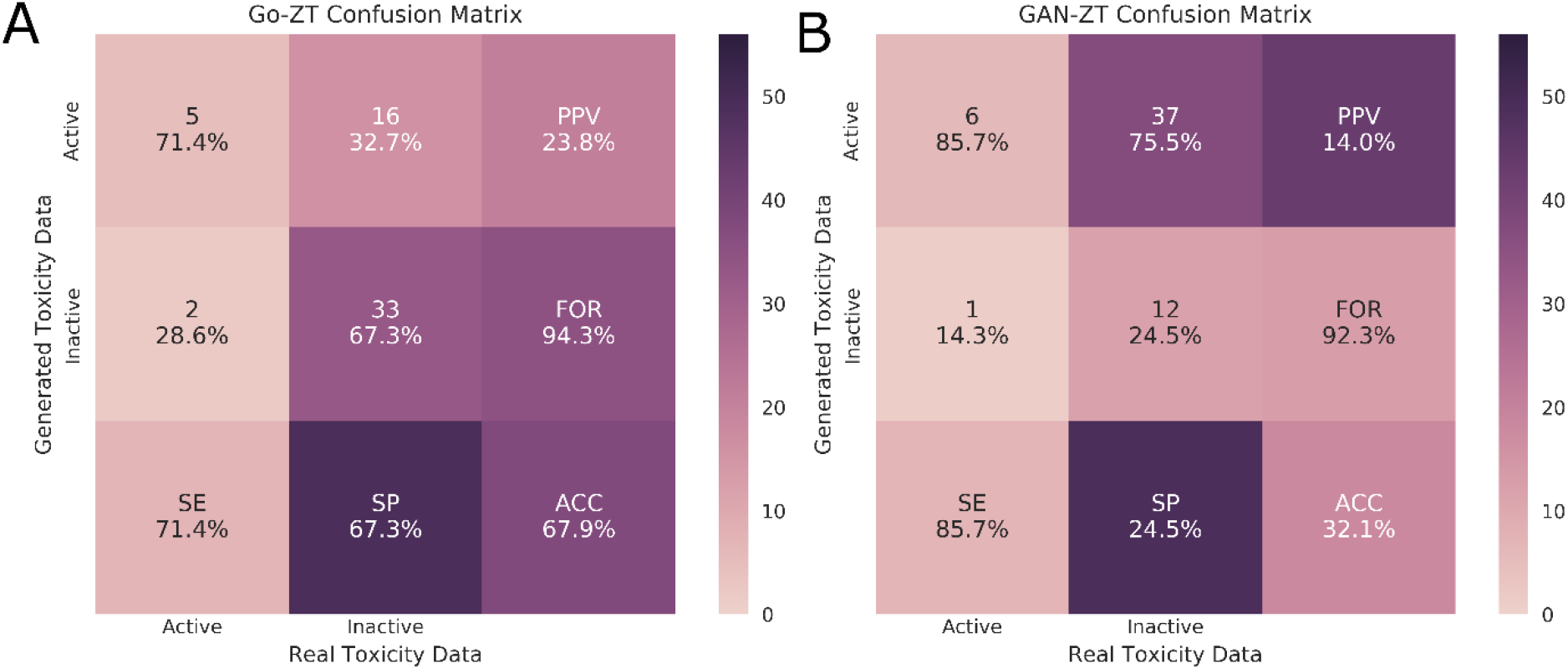
Test dataset confusion matrices. Evaluation of the classification of chemicals in the test data set as either active or inactive using real versus generated toxicity matrices by Go-ZT_v3 (A) or GAN-ZT_v2 (B). PPV – positive predictive value, FOR – false omission rate, SE – sensitivity, SP – specificity, ACC – overall accuracy.

**Fig 6.**
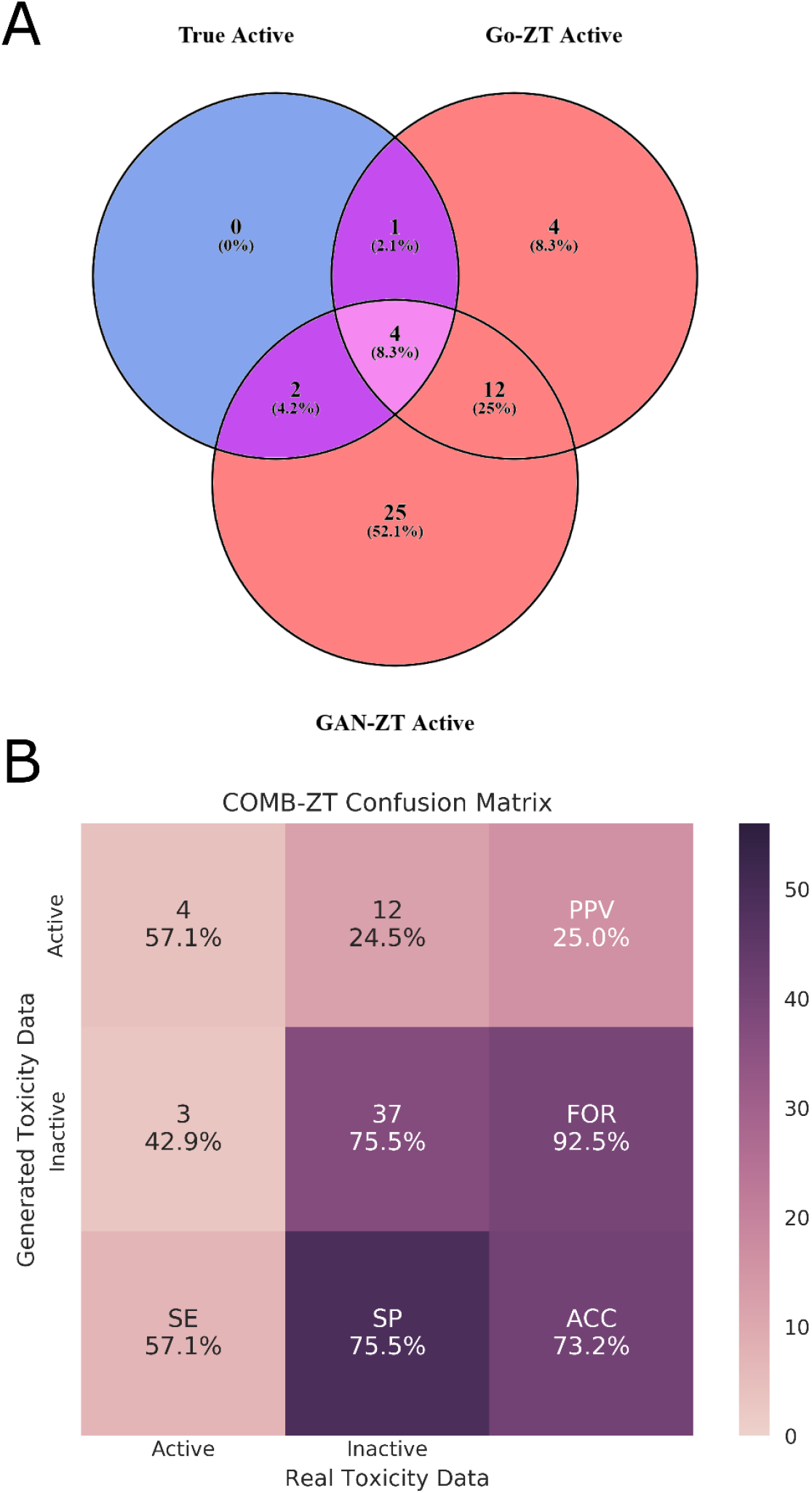
Model consensus on chemical activity. Venn diagram showing the overlap between true active chemicals and chemicals predicted to be active by either Go-ZT_v3 or GAN-ZT_v2 (A). A confusion matrix showing the performance measures of the combined Go-ZT_v3 and GAN-ZT_v2 models (B) using the test dataset.

## Discussion

GAN-ZT and Go-ZT architecture with chemical structure vectorization using views predicts empirical toxicity results with fair Kappa values of 0.125 and 0.149, respectively. Evaluation of an independent and more diverse chemical test set showed similar Kappa, AUROC, and PPV values for Go-ZT while GAN-ZT Kappa, AUROC, and PPV performance declined. Go-ZT_v3 predicted active chemicals with an SE of 71.4%, SP of 67.3%, and a PPV of 23.8% while GAN - ZT_v2 predicted active chemicals with an SE of 85.7%, SP of 24.5%, and a PPV of 14.0%.

When we examined the overlap in predicted active chemicals between Go-ZT and GAN-ZT we see that a consensus model improved the SP, PPV, and Kappa, to 75.5%, 25.0%, and 0.211, respectively but decreased the AUROC slightly to 0.663.

These results show that both supervised generative adversarial and regression-based machine learning architectures are capable of predicting toxic developmental activity with fair to moderate efficacy, especially when we consider the diversity of the test set. Further, leveraging the strengths of both architectures the intersection between the models was able to accurately predict the toxicity of chemicals not part of the initial ToxCast screen.

Other machine learning methods such as Deep Neural Networks (DNN), Support Vector Machines (SVM), *k*-nearest neighbor (*k*-NN), gradient-boosted decision trees, and Bayesian Classifiers have been applied to cheminformatics problems to predict biologically active chemicals with AUROC values ranging from 0.7 – 0.83[31]. These studies utilized ECFP4 fingerprints of chemicals from the ChEMBL database, and single biological activity prediction in drug discovery but were not focused on *in vivo* toxicity. More recent studies have used data from the Tox21 database as part of the NIH 2014 Tox21 Data Challenge with multitask DNNs outperforming other machine learning methods with AUROC values ranging from 0.69 – 0.92 in 12 different biochemical assays[20]. Further, Mansouri and Judson successfully built a QSARs model for G-protein coupled receptor assays using partial least square discriminant analysis that resulted in a balanced accuracy of 96%[32]. Our combined model produced a similar AUROC (0.663) and a lower balanced accuracy value (66%). To the best of our knowledge, this is the first study to develop a DNN model without explicit use of molecular descriptors to predict *in vivo* toxicity in a large chemical set.

While our model is a potentially useful tool for prioritizing chemicals for screening tests, it does have its limitations including high computational costs for training and chemical domain space limits. A better generator could be developed with additional resources. In addition, our networks are dependent on accurate 3D chemical structural information to produce reliable results and at this time are not designed to evaluate mixtures.

Overall, our results show that a DNN utilizing 3D chemical structural information is a useful prescreening tool for predicting the toxic outcomes of the approximately 80,000 untested chemicals registered with the EPA. If we consider that between the training, validation, and test set there are 1,060 chemicals and only 145 are active (13.7%) then there are possibly 10,900 untested active chemicals registered. If the PPV holds at 25% this would result in a list of ~45,000 chemicals to screen. While still a very large number it would reduce the experimental space by over half. Further, these compounds may then be ranked by AggE and the highest chemicals identified should then be prioritized for assessment using the zebrafish HTS assay for developmental toxicity, as these assays are considerably faster and cheaper than traditional chemical screens in mammalian systems.

Looking to the future, increasing computational resources and chemical structural data, alternative network architectures, and inclusion of ToxCast assay results and zebrafish behavioral endpoints in conditional training could improve the predictive value of DNN in *in vivo* toxicity testing. Additional work needs to be done to assess the utility of GAN architecture as a tool to evaluate structure activity relationships (SAR) in toxicology and finally DNNs need to be adapted to evaluate mixtures if a sufficiently large dataset is available.

## Acknowledgements

This research was supported by the National Institutes of Health, through the National Institute of Environmental and Health Sciences (P30 ES030287, R56 ES030007, P30 ES025128) and the National Cancer Institute (R01 CA161608), and the Statistical and Applied Mathematical Sciences Institute.

## Supplemental materials

**S1 Fig.**
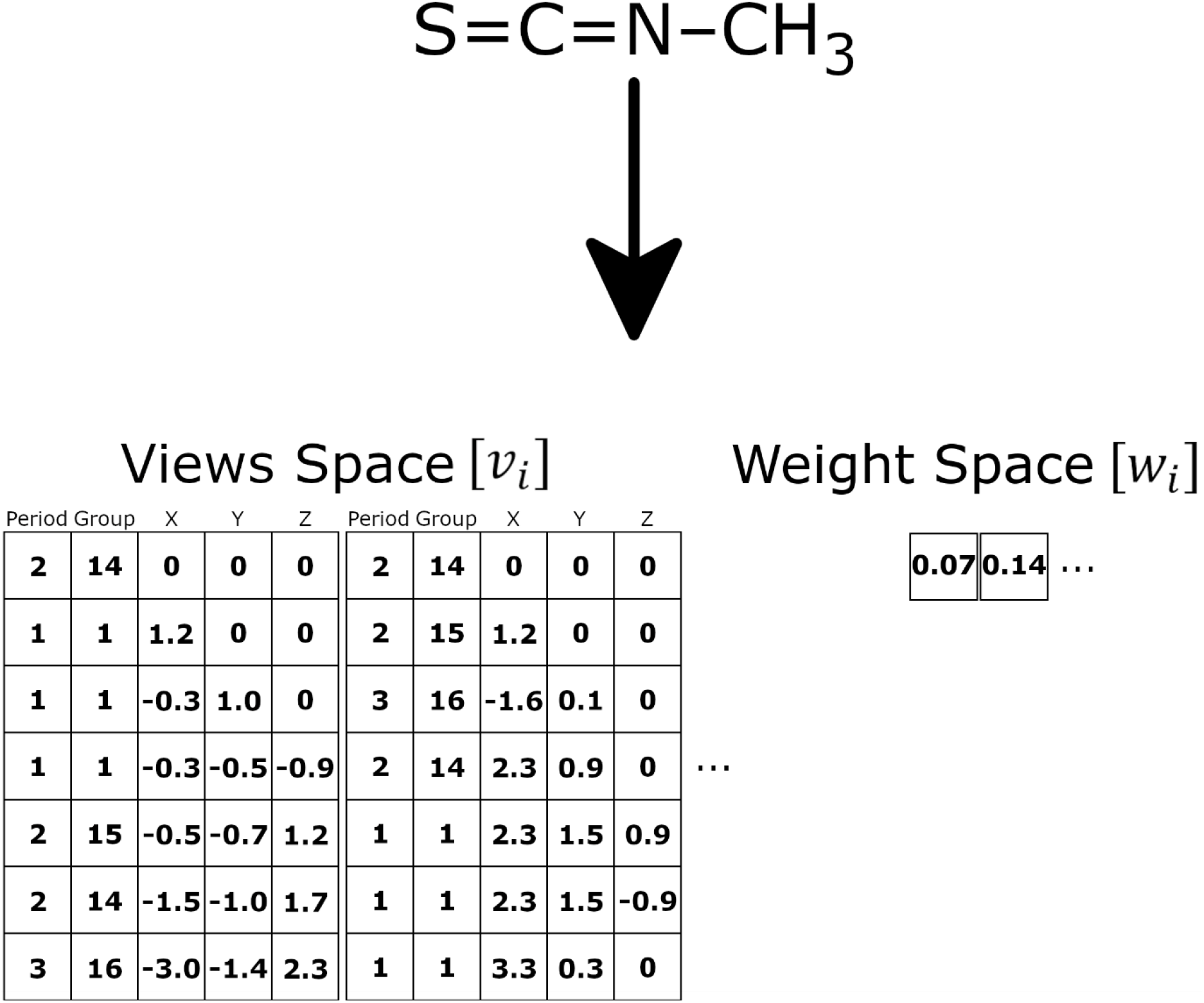
Views vectorization. The views space (vi) columns one and two identify the chemical species and correspond to an atom’s position on the periodic table indicating their period and group, respectively. While the last three columns show the relative position of each atom. The weight space (w_*i*_) values correspond to each of the views space matrices. This molecule has nine views, which can be reduced to three views if preference is given to carbon.

**S2 Fig.**
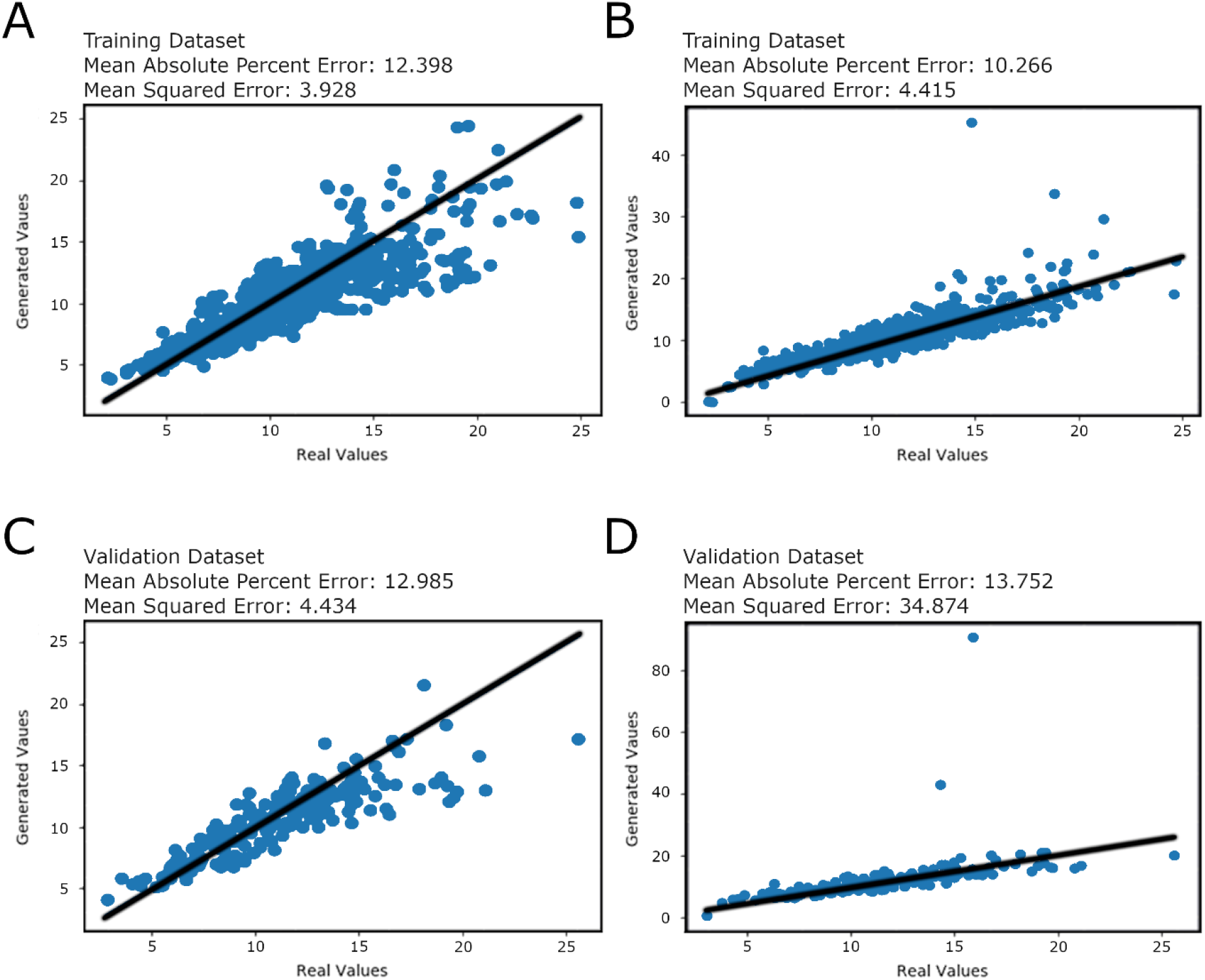
Network efficiency test – generating chemical properties. (A) and (C) shows how well the cGAN architecture (GAN-ZT) generates chemical diameters from the structural data provided by the training and validation datasets. (B) and (D) shows how well the regression architecture (Go-ZT) generates chemical diameters from the structural data provided by the training and validation datasets. Black lines shown are diagonals.

**S3 Fig.**
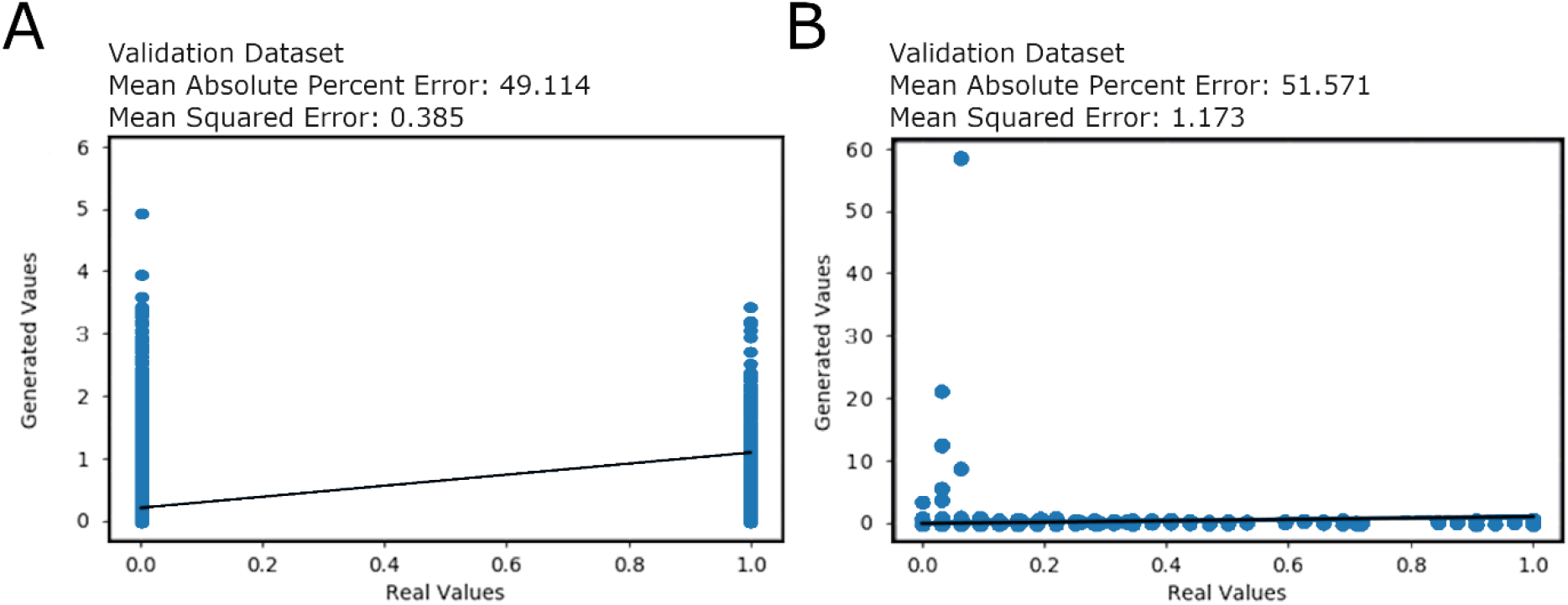
Training results. This figure compares real (empirical) vs generates toxicity endpoint values for the validation chemical set along with summary metrics for (A) GAN-ZT_v3 using individual (binary) toxicity values and (B) Go-ZT using summed and normalized toxicity values.

## Notes

### Competing Interest Statement

The authors have declared no competing interest.

### Summary of Updates

In this version the supplemental materials have been revised and the formatting was updated to reflect requirements for PLOS Comp Bio submission.

